# Optimal inaccuracy: estimating male fitness in the movement-assisted dichogamous species *Clerodendrum infortunatum*

**DOI:** 10.1101/2020.07.25.221127

**Authors:** Amritendu Mukhopadhyay, Suhel Quader

## Abstract

In hermaphroditic species, sexual interference can drive the evolution of dichogamy, where sporophylls (reproductive parts) are separated in time. However, the separation of sporophylls can lead to pollination inaccuracy, especially in movement-assisted dichogamy, where sporophylls alter their position over time. Is pollination inaccuracy minimised by the second sporophyll taking the exact position of the first? Are the sporophylls optimally positioned and stable in their respective active phases? We address these questions in *Clerodendrum infortunatum*, a protandrous, movement-assisted dichogamous species. We made predictions from optimality arguments, and tested these by measuring sporophyll angles over time, by experimentally manipulating flowers, and by estimating correlates of the resultant fitness, taking into account pollen export, pollination inaccuracy and the resultant total pollen delivered. Contrary to expectation, anthers do not have a fixed position in the male phase, and pollination inaccuracy is high. Further, when pollen load is highest, anthers are paradoxically not positioned at the pollen export peak. Also, pollen export and pollination accuracy peaks do not align. This seeming maladaptiveness of anther positioning nevertheless results in highest overall male fitness, measured as the total pollen delivered over the entire male phase. Instead of a simple positional exchange of sporophylls, stamens display a more complicated dynamic strategy which appears close to optimal even though naive measures of pollination inaccuracy are high. Such a strategy of maximising overall male fitness, integrating over the dynamics of stamen trajectory, could well be a general characteristic of protandrous movement-assisted dichogamy.

## Introduction

The diversity of flowering plants is breathtaking. The vast number of species are accompanied by remarkable variation in their life history strategies. One such strategy is the sexual system of flowering plants ^1^, which ranges from monoecy (male and female flowers occur on the same individual) to dioecy (male and female flowers occur on different individuals), with several variations, including androdioecy, andromonoecy, and many more. The most common sexual strategy in flowering plants is hermaphroditism (male and female sexual organs occur in the same flower, and the plant bears only this type of flower), which occurs in nearly 72% of all angiosperms ^2^.

Why should hermaphroditism be so common in flowering plants? The theory of plant resource allocation in economic terms ^3,4^ suggests that this is the most efficient strategy as it minimises resource investment. This is because a common investment in attractive traits, including petals and nectar, promotes both male and female reproductive success at the same time. However, this advantage comes with an associated cost – that of self-pollen deposition. This cost is borne by both self-compatible and self-incompatible species. In self-compatible species, self-pollen deposition increases the possibility of inbreeding depression, where offspring vigour decreases down the generations ^5,6^. In addition, primarily male reproductive success might decline because of sexual interference ^7–9^. Sexual interference leads to wastage of pollen because deposited self-pollen on stigmas cannot be deposited on stigmas of other individuals and are therefore removed from the cross-pollen pool. On the other hand, in self-incompatible species, sexual interference is the only cost to bear.

Flowers have evolved different strategies to minimise these costs. The details vary from species to species, but the general theme is either spatial or temporal separation of sporophylls (stamen and pistil) in the flower ^7,10^. In herkogamy, both sporophylls are active (reproductively mature) at the same time, but separated in space. By contrast, in dichogamy, sporophylls become active at different times. In this way, dichogamous flowers have separate male and female phases. In the special case of movement-assisted dichogamy, the sporophyll that matures earliest is positioned in the presentation zone (where it exports or receives pollen, as the case may be) in the first phase. In the second phase, it moves away and the second sporophyll becomes active and takes that position.

Any spatial separation of sporophylls has an associated problem of pollination inaccuracy ^7,11^. For successful cross-pollination, the pollen export and receipt positions must be similar among flowers in a population. This is because, in vector-pollinated plants, pollen transfer is maximum when the part of the pollinator’s body that touches (and therefore picks up) pollen from the anthers is the same part of the body that touches (and therefore deposits) pollen to the stigma. Because particular species of pollinators tend to show stereotypical foraging postures at flowers, this requirement is most closely met when anthers and stigmas in different flowers are at exactly the same position. For this reason, any spatial separation between anther and stigma necessarily increases pollination inaccuracy even as it serves the purpose of decreasing self-pollen deposition.

Pollination accuracy or inaccuracy, then, clearly affects optimal presentation of the sporophylls. A reasonable starting point would be that the optimal position of a mature sporophyll is where the sporophylls has the highest probability of touching the relevant body part of the pollinators; this leads to the highest pollen export and deposition. It follows from this that mature sporophylls in natural populations should occupy this ‘optimal presentation zone’. Though early natural history studies ^12–14^ took this for granted, the assumption is rarely tested explicitly ^11^.

Different floral adaptations lead to varied consequences. The permanent spatial separation between sporophylls of herkogamous species is expected to lead to high pollination inaccuracy ^8^. In dichogamous species, by contrast, the first sporophyll makes way for the second, and it is potentially possible for both sporophylls to occupy the same position, albeit separated in time. Does this mean that pollination inaccuracy is lower in dichogamous species than in herkogamous species? Unfortunately, few studies have attempted to answer this question ^15^. Species that show movement-assisted dichogamy present a further complication as their sporophylls are in motion. In this case, the theory of optimal sporophyll placement depends on an additional assumption that after entering the optimal presentation zone, mature sporophylls remain stationary. However, this is not always the case ^16^. If sporophylls are not stationary in the presentation zone, then pollen export and deposition will vary dynamically as the sporophylls move. This means that fitness of traits needs to be estimated across the lifetime (and therefore position) of sporophylls to understand the overall outcome.

Therefore, a detailed study on adaptive fitness of movement-assisted dichogamy should address the following aspects in sequential order. First, whether male and female sporophylls take on the same position during maturity. Second, whether they both take on ‘optimal’ positions. To assess that one needs to compare indices of fitness between the observed phenotype with other variants ^11,15,17^. These variants can come from among the natural variation within populations, but when that variation is small (possibly due to strong selection), artificial variants must be created. Morphological trait manipulations are routinely used by floral biologists to create artificial variants ^18–20^. Even, within the range of natural variation, one may need to create artificial variants in order to study dynamic traits (e.g., sporophyll position in movement-assisted dichogamy). To estimate a fitness index of a sporophyll at a specific position, one needs to create an artificial variant, which is fixed at that position over time.

A population may depart from optimality (called adaptive inaccuracy) in one or both of two ways: the mean of the population lies away from the optimum (called bias), and phenotypic variation is high such that many individuals are far from the optimum (called imprecision) ^21^. In a perfect world, with all members of the population perched on the adaptive peak, both bias and imprecision are zero. However, maladaptation in the population can reveal itself through nonzero bias, or nonzero imprecision, or both. Depending on the situation, selection may be directional, or stabilizing, or both.

To investigate the adaptive fitness of movement-assisted dichogamy in detail, we studied the complete (no overlap between sexual phases) dichogamous species *Clerodendrum infortunatum* (henceforth *Clerodendrum)* ^16^. Though *Clerodendrum* is a self-compatible species, it mainly breeds by xenogamy and geitonogamy ^22,23^. Thus, for reproduction, *Clerodendrum* in natural population heavily depends on pollinators. We predicted that pollination inaccuracy of the natural population is low. We also tested the prediction that in flowers, anthers and stigma are positioned optimally to maximise their fitness. In manipulated flowers, we fixed anthers and stigma at various positions within and beyond their natural ranges, and estimated pollen export and deposition rate at each position. We predicted that the resultant curves would be unimodal, where the highest peaks indicate the optimal presentation zone for each of the sporophylls. We expected the peaks for anthers (pollen export) and stigmas (pollen deposition) to overlap strongly. We then investigated whether anther and stigma positions in the natural population overlap with the expected male or female optimal presentation zones as estimated through the morphological manipulation experiments. We estimated cumulative male fitness over the entire male phase. We also estimated bias and imprecision as a measure of the degree of maladaptation in the natural population.

## Methods

### Observations and experiments

The Anamalai Hills, dominated by mid-elevation tropical wet evergreen rainforest ^24^, are situated in the southern Western Ghats in India. The Valparai plateau (700m-1600m asl) lies within these hills. Though the Valparai landscape is dominated with tea and coffee plantations, many privately owned rainforest fragments (ranging from 0.3 to 650 ha in size) still persist. These forest fragments harbour a significant diversity of rain-forest flora and fauna ^25,26^. Two such forest fragments were selected for this study, namely Iyerpadi Riverine (10°21’12“ N, 76°59’54” E, Area 1.76 ha) and Varattuparai Colony (10°21’22“ N, 76°56’30” E, Area 33.4 ha). Within these forest fragments, we studied the shrub *Clerodendrum infortunatum* Gaertn. [Lamiales: Lamiaceae] between December 2014 and May 2015. *Clerodendrum* produces dichogamous and protandrous flowers, where the male phase precedes the female phase. Flowers are mainly pollinated by butterflies ^22,23^.

To describe the spatiotemporal dynamics of sporophylls, we monitored flowers between 18 February 2015 and 26 April 2015. Thirty two flowers from 18 plants were tagged with coloured thread, so that they could be monitored throughout their lifetime of two or more days ^16^. We monitored flowers every two hours from the morning (6 AM) to evening (6 PM), with a total of seven visits per day. The next visit was on the morning of the next day. During each visit, a lateral photograph of the flower was taken. Stamen and style angles were measured later from the photographs using ImageJ software (Fiji distribution) ^27^. In order to measure sporophyll positions in a consistent manner, we defined the geometry of a flower as follows. Taking the horizontal floral axis as the starting point, the basal direction of the plane was defined as being at an angle of 0° and, measuring counter-clockwise, the distal direction as 180° (Fig. 1). Angles from this plane to the anthers and stigma were measured from the point where the floral axis was intersected by an imaginary vertical line along the plane where corolla tube opens up.

**Fig. 1.**
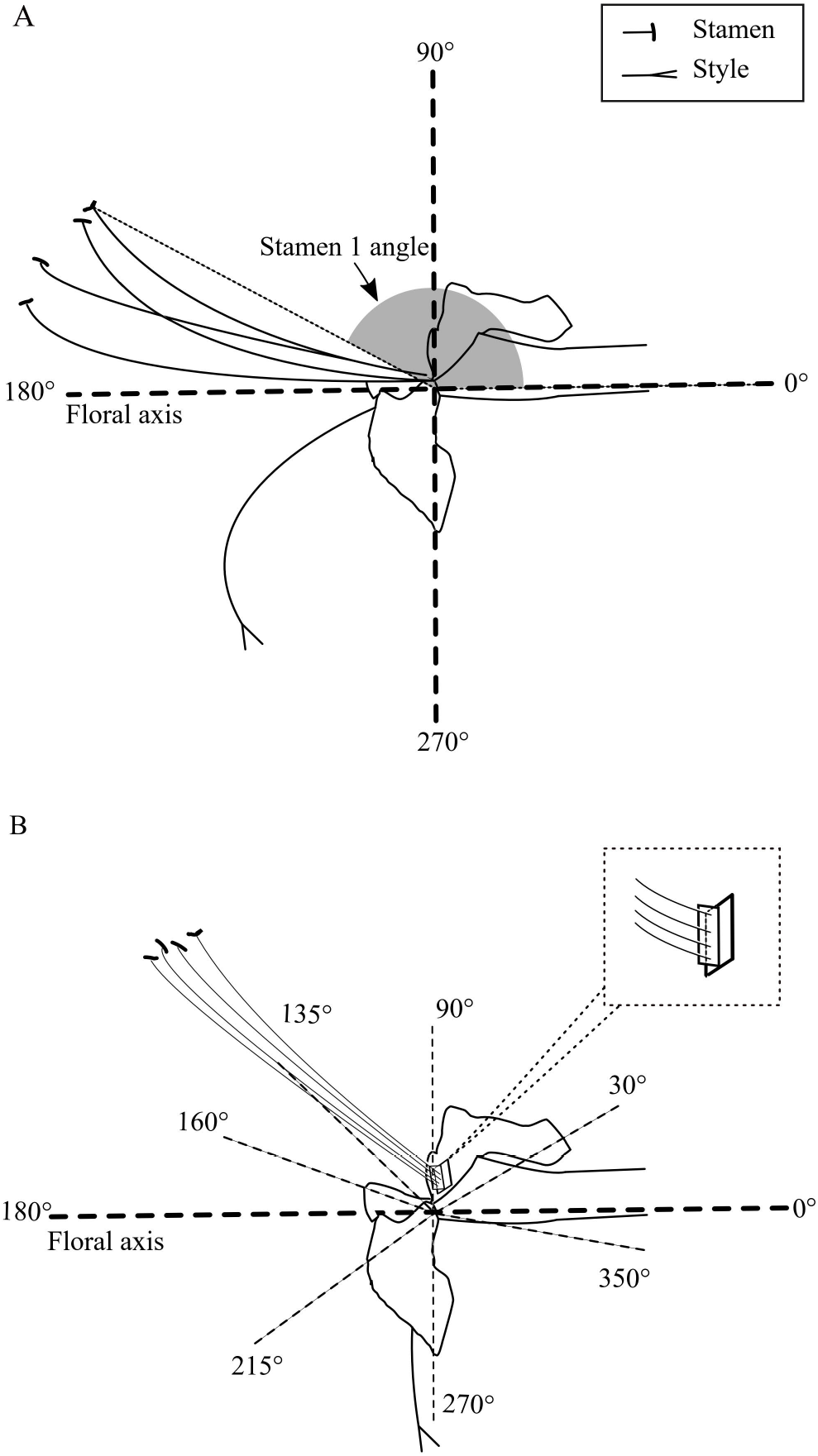
A. A natural *Clerodendrum* flower at male phase. B. A schematic diagram of a morphologically manipulated flower, where anthers were set at a fixed position near 135°. All angles [30°, 90°, 135°, 160°, 215°, 270°, 350°] at which sporophylls were set in different treatments are also shown.

The fitness consequences of adopting different phenotypes (sporophylls at different angles) were measured by experimentally manipulating floral structures. Because sporophylls are in continuous motion, it is impossible to find sporophylls at a fixed position in natural flowers. Also, sporophylls take on a narrow range of positions in their active phases, and the consequences of adopting a position outside this range cannot be assessed in natural populations. Flowers were therefore manipulated to set the active sporophyll at a specific angle, and the other sporophyll at its normal inactive position (roughly 270°) (Fig. 1). These experiments were carried out between 28 April 2015 and 24 May 2015. All morphological manipulations were done on the male stage flower (target flower) so that nectar content was roughly similar and could not influence visitation differentially. *Clerodendrum* flowers are white, and so white double-sided adhesive tape was used to attach the sporophylls (after removing the existing sporophylls) to the flower at a specific angle. The tape was without any detectable odour, which might alter visitor behaviour. On a small piece of tape, stamens or style were attached and then that piece was attached to another small piece at 90° (Fig. 1). This second piece was attached to the flower. Stamens and styles used in this experiment were collected from a separately bagged inflorescence to make sure that the used stigmas did not contain any pre-deposited pollen, and the pollen loads of the used anthers were intact. Styles were collected from female-phase flowers and stamens were collected from male-phase flowers in the donor inflorescences.

In this manner, active sporophylls were experimentally set at the following angles – 30°, 90°, 135°, 160°, 215°, 270° and 350°. These angles were chosen in order to cover virtually all possible approach angles around the flower. Treatments in which sporophylls were affixed towards the distal end of the flower (< 90° and > 270°) were unlikely to receive pollen from pollinators, but provide an estimate of pollen deposition from other sources (e.g., from the shaking of branches). To check for possible adverse effects of manipulation, we compared sporophylls manipulated to be in the presentation zone with unmanipulated sporophylls (see Results section). Pollen of unmanipulated flowers was quickly depleted (personal observations), and hence flowers with manipulated stamens were kept exposed for 6 hours (0800 to 1400 hrs). By contrast, pollen deposition rates were low, so flowers with manipulated stigmas were kept exposed for 24 hours (0800 of day 1 to 0800 of day 2). The exposure durations were about 3/4th and 3/5th of the natural male and female phase durations respectively ^16^. We have assumed that pollen export and deposition rate is zero at 0° and 359° (the distal end of the flower), since in nature, the pedicel blocks the pollinator approach path at those angles. We also assumed that pollen export and pollen deposition rates over the duration of the experiments (6hr and 24hr respectively) were constant for a particular sporophyll position. After the exposure period, sporophylls were collected in vials containing 70% ethanol: all four anthers from a flower in a single vial and the stigma in a separate vial. This storage method has a caveat – pollen grains loosely attached to the stigma surface might get washed off. However, the remaining pollen on the stigma is likely to provide a reasonable estimate of the number of germinating pollen since germinating pollen adheres tightly to the stigma surface because of pollen-pistil interaction ^28,29^.

Pollen were counted in the lab, from 28 April to 21 June 2016. Stigmas were excised and placed on microscope slides; they were first washed in 95% ethanol, and then stained with Safranin O ^30–32^. After a drop of glycerine was added, the stigma was squashed under a coverslip, and pollen were counted under a compound microscope (100x magnification). Because *Clerodendrum* anthers have a large number of pollen grains, it was not possible to conduct a total count of the leftover pollen on anthers, and so a suitable sampling method needed to be devised. We used a slightly modified version of the protocol described by Shivanna ^33^. Anthers were placed in a 1 mL vial with 70% ethanol solution. A drop of detergent was added to keep the pollen suspended. The anther was crushed with a glass rod to release pollen grains. The suspension was mixed thoroughly by repeatedly drawing it into and expelling it from a syringe without the needle. While gently shaking the vial we took a 0.01 mL volume of solution with a micropipette. The solution was spread on a haemocytometer glass slide, covered with a coverslip. The total number of pollen grains in the solution was estimated by averaging the number of pollen grains across the ten replicate subsamples, dividing the average by the volume of the subsample (0.01 mL) and then multiplying it by the total suspension volume (1 mL).

### Analyses

All angular data were summarised using circular statistics. Because anther and stigma angles in active phases (i.e., male and female phases respectively) were in a small range (between 150° and 180°), simple linear differences were calculated where needed. We arrived at an operational definition for determining male and female phases by defining an angle of transition between these two floral phases. This transition angle was set as 180° (based on sporophyll movement ^16^). When the average anther angle was less than (greater than) the critical anther angle, the flower was considered to be in the male phase (female phase). The critical angle is also the transition angle, so when the average anther angle is less than 180°, this implies that the style angle is greater than 180°. In this way, although we have defined phases based only on average anther angle, the definition captures the position and movement of the stigma as well. At anthesis (hour 0), a large proportion of the anthers remains undehisced, and stamens remain partly coiled (personal observations). For this reason we consider the initial anther stage to begin at hour 2, when pollen from the dehisced anthers become available for export.

Positional mismatch, which can lead to pollination inaccuracy, is estimated as the absolute difference in measurements between the mean anther angle, and the mean stigma angle of the natural population in their respective active phases. As the average anther angle changed with time in the male phase (Supplementary Fig. 1), positional mismatch was estimated at different time points in the male phase.

Using the experimental constructs, which manipulated sporophyll position, we estimated pollen removal (from anthers) and pollen deposition (to stigmas), and with this, modelled male fitness in relation to sporophyll angle. A phenomenological modelling approach was used, where the best fit model was used to describe the relation, and parameter values were either estimated from the data (see Results section) or collected from the literature.

The number of exported pollen cannot by itself be used as a measure of male fitness. Pollen export by vectors is very inefficient process, where most of the exported pollen does not deposit on conspecific stigmas ^34^. A better measure of male fitness is the estimated number of exported pollen that is eventually deposited on conspecific stigmas. This depends on a combination of pollen load, export rate, and the proportion of exported pollen that is deposited per unit time (deposition rate). Following from the theory of pollination accuracy, we assumed that pollen removed by pollinators from an anther at a particular position (angle) is deposited on stigmas at that same position. In all, the male fitness of a flower with anthers at a specific angle is a combination of the pollen export rate, pollen deposition rate and available pollen load on anther at that angle. For the purposes of this study, we assumed that all pollen deposited on a flower comes entirely from another single flower. If this assumption is violated, the pollination efficiency parameter would change; we explore the possibility in the Results section.

Pollen export rate was modelled using results from the morphological manipulation experiment. For each manipulated treatment, we subtracted pollen counts of the anthers (retained pollen) from the average initial pollen load of *Clerodendrum*, which is 6082 ^22,23^. The proportion of pollen exported from specific anther positions was calculated per 0.36 hr, which is the time taken by anthers to move 1° (Supplementary Fig. 1). Pollen export rate is not constant across the male phase, but rather changes with anther position. Export rate is initially low, and as anthers move towards the floral axis, the rate increases and then decreases. Pollen export rate *E* was modelled as a function of anther angle (*x*) in the following manner (Eq. 1). Parameter *n* is the anther angle (on a linear scale) where pollen export rate would become 0. Parameter *m* (*m* > 0) is the visitation rate. With the increase in number of visits pollen export rate increases.

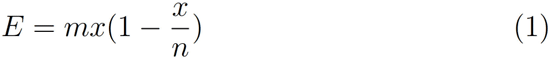

The number of pollen deposited at any specific angle was divided by the estimated number of pollen exported from that angle to estimate the proportion of exported pollen that was deposited on a stigma at that specific angle. The proportion of exported pollen that was deposited per 0.36 hr (the deposition rate curve) (A) was modelled as a function of anther angle (*x*) using a bell-curve function with a single peak (Eq. 2). Parameter *a* (*a* > 0) represents the height of the curve, which is the indicator of the efficiency of the pollinator. Parameter *b* is the angle at which pollen deposition is highest. This angle corresponds to the approach angle of the pollinators. Parameter *c* (*c* > 0) is the strength of pollination accuracy selection. With increasing strength of pollination accuracy selection, the pollen deposition rate curve becomes narrower (Supplementary Fig. 2).

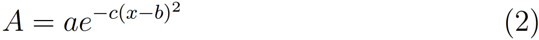

Pollen delivery rate depends on the pollen export rate and exported pollen’s rate of deposition onto conspecific stigmas. The deposition rate depends on pollination accuracy. Therefore, the pollen export rate function (Eq. 1) was multiplied by the pollen deposition rate function (Eq. 2) to estimate pollen delivery rate (Eq. 3) with respect to anther angle. This equation estimates the proportion of exported pollen delivered to conspecific stigmas per unit time (0.36 hr) at a specific anther angle.

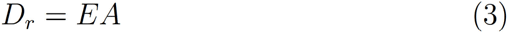

The available pollen load (*P*) on anthers changes with time as stamens move. Available pollen depends on the initial pollen load (*P(v)* = 6082, see above) and pollen export rate (*E,* Eq. 1). The change in pollen load on anthers with time can be described by an extension of the exponential population growth equation, where the rate is negative and depends on anther position. The anther angle (*x*) was used as a proxy for time, as it changes with time in a linear fashion for most of the duration of the male phase (Supplementary Fig. 1). Thus, anther pollen load is modelled here as a function of anther angle rather than as a function of time. Change in pollen load (*dP*) with change in angle (*dx*) depends on export rate *j* and pollen load *P*. The rate parameter *j* was replaced by the Eq. 1. By integrating both side (definite integration, where *v* is the initial anther position and *s* is the anther position, where pollen load needs to be calculated) we derive an equation (Eq. 4) that describes pollen load at different anther positions (*P(s)*).

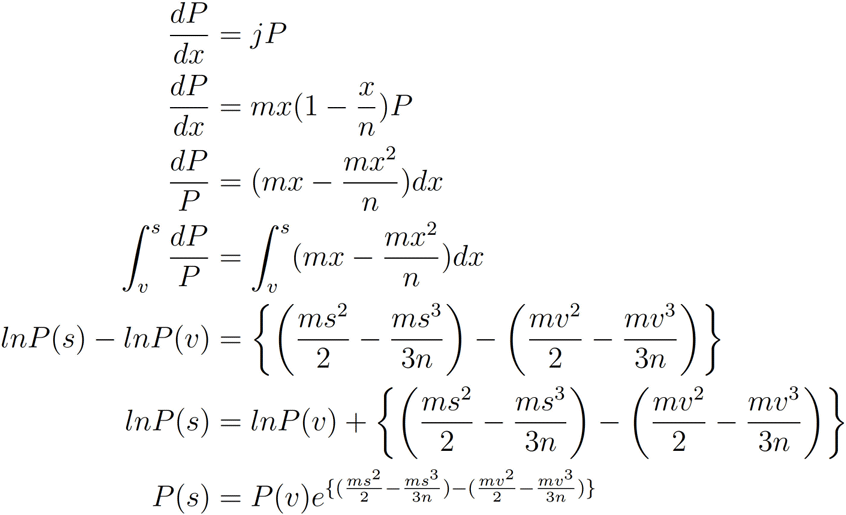

Because the rate is negative (pollen load decreases with time as anthers move) the exponent in Eq. 4 is negative.

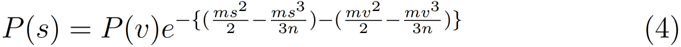

The proportion of pollen delivered to conspecific stigmas per 0.36 hr at different anther positions is described by Eq. 5. This is the male fitness curve, and is the product of available pollen load (Eq. 4) and the pollen delivery rate (Eq. 3).

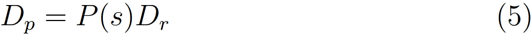

As active anthers are not stationary, overall male fitness (Eq. 6, total pollen delivered to conspecific stigmas in the entire male phase) can be estimated only by summing fitness over the lifetime of the anther (i.e., over the multiple positions it takes). This was done by integrating the resulting male fitness function between the limits *v* (anther starting angle, when anthers reach maturity) until *u* (180°– the transition angle, when sexual interferences take place ^16^). We have assumed that sexual interference is the key force driving the evolution of dichogamy in *Clerodendrum*. A model has shown that sexual interference alone can lead to the evolution of dichogamy ^35^. Also, as flowering is asynchronous in *Clerodendrum* and geitonogamy takes place ^22,23^, dichogamy can not help flowers much in reducing selfing. However, it can minimise the sexual interference.

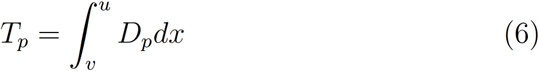

After the male fitness curve is estimated, it becomes possible to estimate the bias and imprecision in the degree to which natural phenotypes match the optimum (the peak of the fitness curve; found by numerical exploration). Bias is defined as the deviation between the fitness of mean of the population and the theoretical maximum fitness. Imprecision is the population variance of fitness, which is the scatter of individuals around the population mean. Adaptive inaccuracy is estimated ^21,36,37^ as the sum of imprecision and the square of bias, with the unit being the square of total pollen delivered.

All analyses were carried out in R, version 3.5.1 ^38^; with the following additional packages: nlme ^39^, boot ^40,41^ and circular ^42,43^. Error bars in figures are 95% nonparametric bootstrap confidence limits. This method circumvents restrictive assumptions about the data distribution ^44^. We draw inferences from the data based on the estimated effects and their uncertainty as measured by their confidence intervals. In addition to the estimation approach ^45–47^, we also presented results of null hypothesis significance tests in which the p-values assess our ability to discern experimental effects statistically.

## Results

In the male phase, the anthers are tightly clustered around 150° (mean vector angle, α = 150.85°; [95% CI = 145.92°, 156.04°]; mean vector length, r = 0.98; N = 24). Similarly, in the female phase, stigmas are closely arranged around 167° (mean vector angle, α = 167.17°, [95% CI = 163.82°, 170.74°]; mean vector length, r = 0.99; N = 24). Although the difference between the mean orientation of male and female sporophylls is only 17°, the low within-group variation means that the difference is statistically discernable (Fig. 2; p < 0.001; Watson-Williams test for homogeneity of means).

**Fig. 2.**
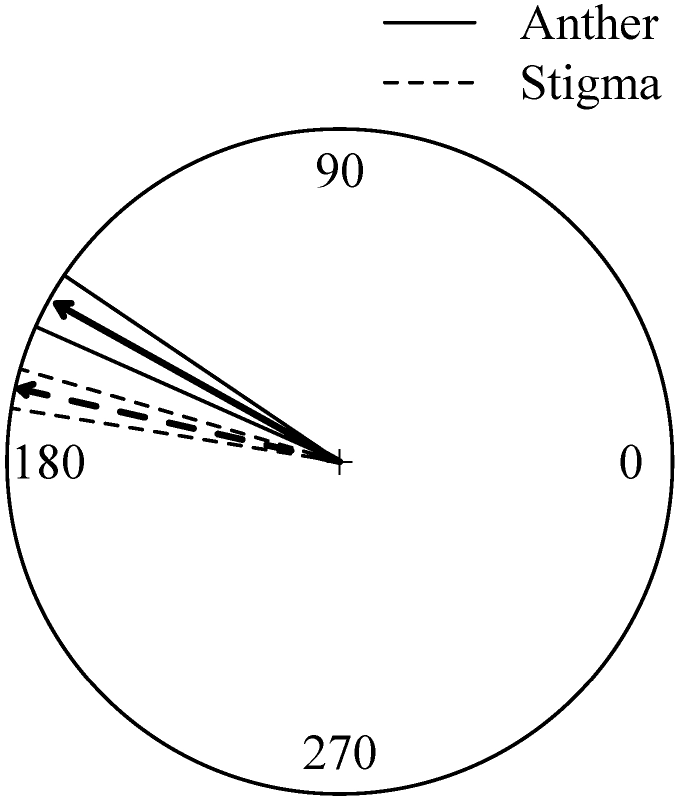
Arrows show circular summary statistics of the angles of anthers at hr 2 in their active phase and the angles of stigmas in their active phase. The direction indicates the mean, and the length signifies the concentration. Radial lines flanking the group mean vectors mark the 95% confidence intervals.

While the stigma remains stable over its entire active phase the anthers move continuously ^16^. For the most part of the male phase, the anther angle changes approximately linearly with time (Supplementary Fig. 1). To be specific, anthers take roughly 0.36 hr to move 1°. The difference in movements between stigma and anther means that the positional mismatch between the sporophylls changes with time. At eight hr from anthesis, positional mismatch is minimal (Supplementary Fig. 3). Subsequently, it increases dramatically towards the end of the male phase.

We compared pollen retention between control (male phase natural flower) and treatment (160° angle floral constructs of anthers) flowers to estimate the effectiveness of manipulated male flowers. Similarly, we compared pollen deposition between control (female phase natural flower) and treatment (160° angle floral constructs of stigma) to estimate the effectiveness of manipulated female flowers. There is a relatively small (and not statistically discernible) difference between retained pollen number in control and treatment (M_control_ = 1203, 95% CI = [768.03, 1730.97], M_treatment_ = 1353.85, 95% CI = [944.62, 1812.23], Welch two sample t-test, t = −0.43124, df = 19.728, p = 0.671, N_control_ = 10, N_treatment_ = 13). Similarly, there is a small (in absolute terms, and not statistically discernible) difference between deposited pollen number in control and treatment, though there is considerable variation (M_control_ = 2.25, 95% CI = [1.12, 3.5], M_treatment_ = 4.2, 95% CI = [1.5, 7.5], Welch two sample t-test, t = −1.1314, df = 11.794, p = 0.2804, N_control_ = 8, N_treatment_ = 10).

As expected, pollen retention for anthers affixed towards the basal end of the flower is high (Fig. 3A), and when Eq. 1, (with parameters *m* = 0.0004300188; *n* = 381.4351) is fitted to the proportion of pollen exported per 0.36 hr, the curve is rather flat at the peak indicating a relatively broad range of positions over which pollen are removed (Fig. 3C). By contrast, the pollen deposition curve (Eq. 2, *a* = 0.00001725803; *b* = 170.0948; *c* = 0.0008193516) shows a sharp peak with respect to position (Fig. 3D).

**Fig. 3.**
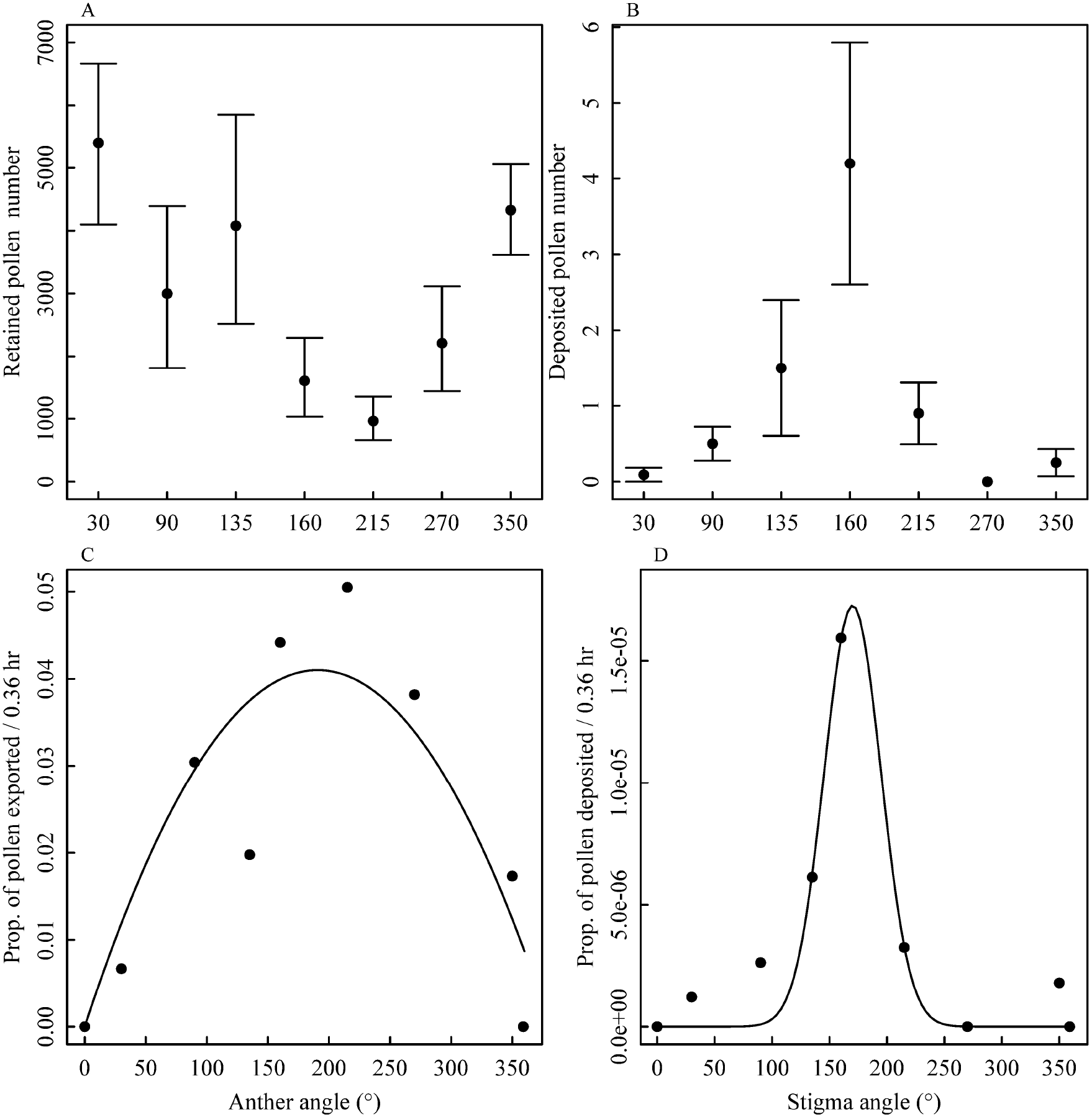
A. Pollen retained at anthers set at specific angles in morphologically manipulated *Clerodendrum* flowers. B. Deposited pollen at stigmas set at specific angles in morphologically manipulated flowers. C. Proportion of pollen exported per 0.36 hr from anthers set at specific angles. The fitted curve representing pollen export rate function (Eq. 1) is plotted. D. Proportion of exported pollen deposited per 0.36 hr at stigmas set at specific angles. The fitted curve representing pollen deposition rate function (Eq. 2) is plotted. Error bars are 95% nonparametric confidence intervals.

When overlaying the pollen export and pollen deposition functions (Fig. 4A), we see that their peaks (190.7° and 170.1° respectively) are separated. The product of these two functions, which describes pollen delivery rate, peaks at 170.8°, virtually identical to the pollen deposition peak (Fig. 4B) because the pollen export function is relatively flat in this region (Fig. 4A). The mean anther angle of the population (150.9°) at the beginning of the male phase (hr 2) turns out to be far away from the delivery peak (Fig. 4B). Available pollen load is highest at hr 2 (150.9°), after which it decreases rapidly (Eq. 4, Fig. 4B). When anthers come to the delivery peak, their pollen has been substantially depleted (Fig. 4B). From the starting position (150.9°) as anthers move, the number of pollen delivered from different positions declines steadily (Eq. 5, Fig. 4C).

**Fig. 4.**
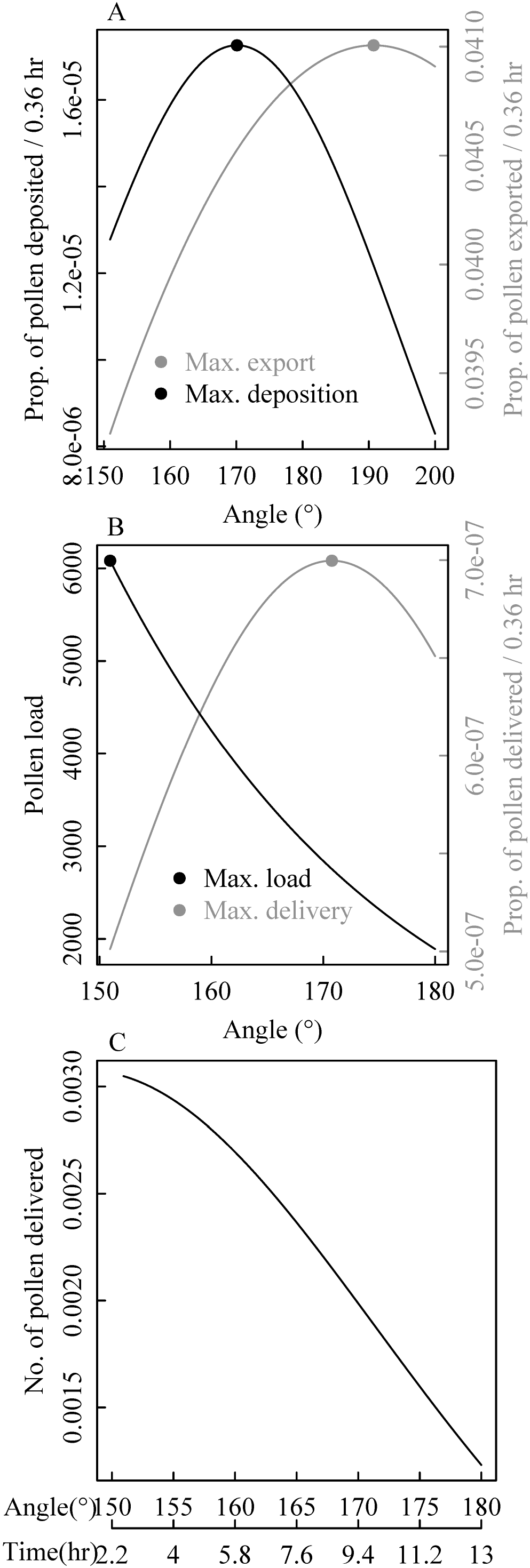
A. Pollen export rate and pollen deposition rate focused on a narrower range of angles. The resultant delivery rate is the product of the two equations. Maxima of each function are indicated. B. Pollen load (Eq. 4) and delivery rate functions in relation to anther angle, with the maxima of each function. C. The resultant delivered pollen at different anther angle (Eq. 5).

The different starting positions of anthers lead to different numbers of total pollen delivered (Eq. 6) to a single stigma from a single set of conspecific anthers (i.e., on a single flower) in the entire male phase (Fig. 5). The highest total pollen delivery would happen if anthers started at 146.3°. Surprisingly, starting at the position of the maximum delivery rate (170.8°) leads to the lowest total pollen delivered. The total pollen delivered in case of the natural starting position (150.9°) of anthers is close to the maximum peak. The overall adaptive inaccuracy of the natural population is 0.0004341271 pollen^2^, with imprecision (0.0004330152 pollen^2^) making a much larger contribution than bias (0.000001111884 pollen^2^).

**Fig. 5.**
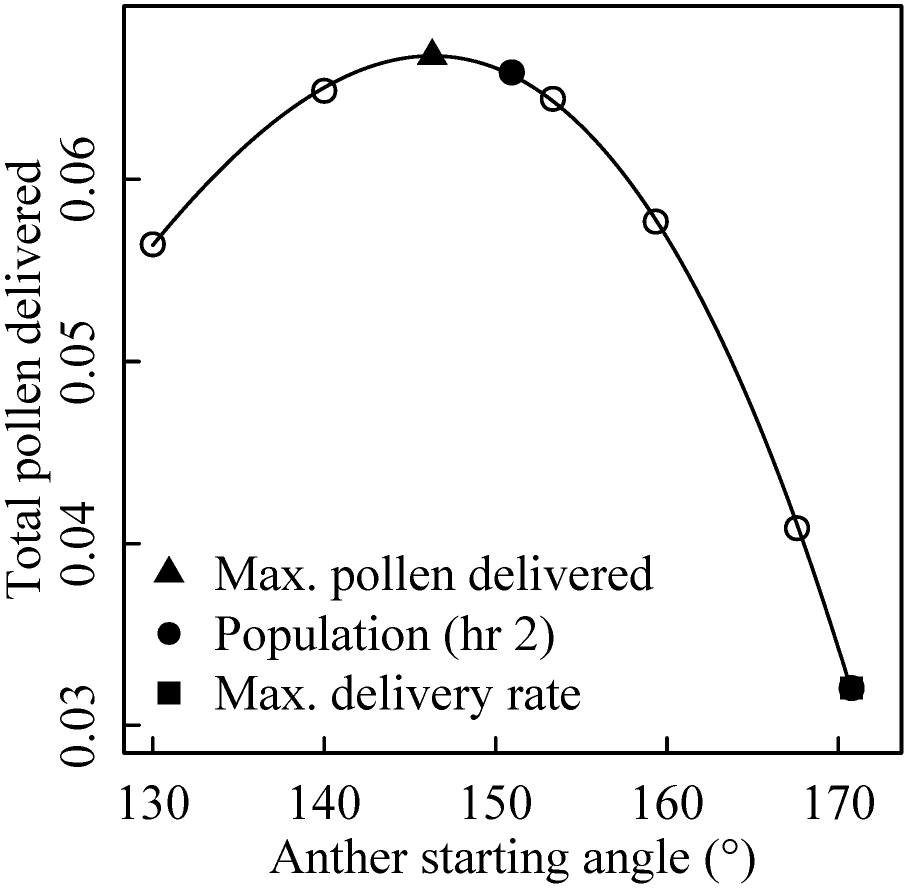
Total pollen delivered (Eq. 6) to a single stigma from a single set of conspecific anthers (single flower) in the entire male phase for different starting positions of anthers. Cubic polynomial function (*y* = – 0.00000047066*x*^*3*^ + 0.0001597981*x*^*2*^ – 0.01653496*x* + 0.5393821) is fitted to the data, where *y* is the index of total pollen delivered in the entire male phase and *x* is the starting position of anthers. The points that indicate the maximum total pollen delivered, total pollen delivered in natural population and total pollen delivered if anthers would start at delivery peak are shown.

With the increase in visitation rate, both the total number of delivered pollen and optimum start angle increases (Fig. 6A). With increasing pollinator efficiency, the total number of delivered pollen increases (Fig. 6B). However, the position of optimum remains the same. By contrast, with increasing strength of pollination accuracy selection, the optimal peak moves closer towards the pollen delivery peak, but the number of total pollen delivered decreases (Fig. 6C).

**Fig. 6.**
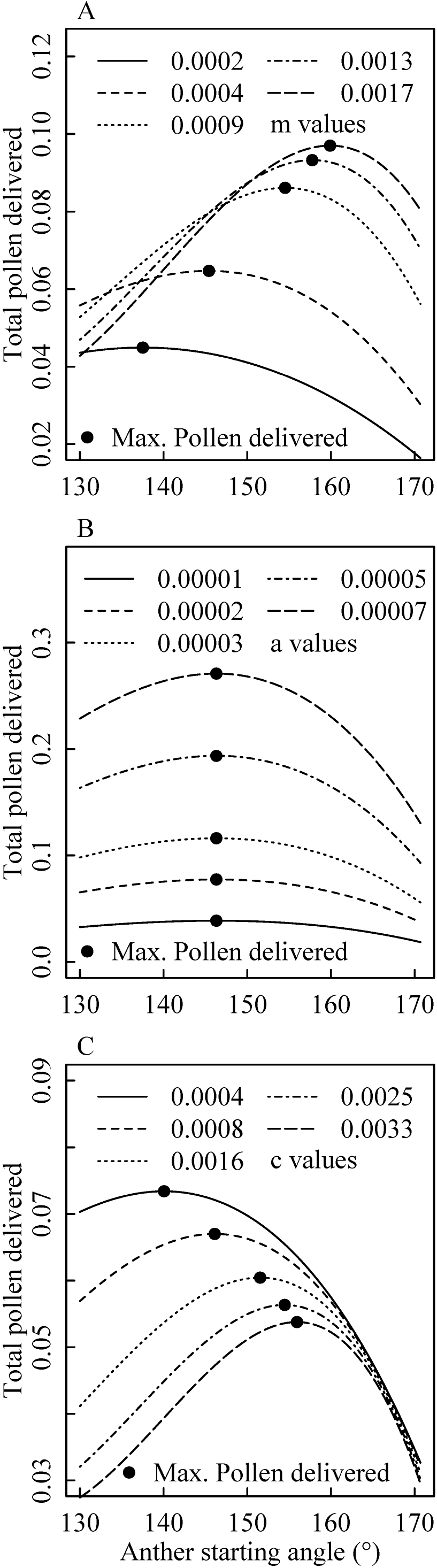
A. The relation among visitation rate (parameter *m*) and number of total pollen delivered (Eq. 6) and optimum starting position of anthers. B. The relation between pollinator efficiency (parameter *a*) and the number of total pollen delivered (Eq. 6). C. Anther starting positions where maximum pollen delivery (Eq. 6) occurs at different strength of pollination accuracy selection (parameter *c*). Cubic polynomial functions are fitted to the data.

## Discussion

Contrary to expectation, *Clerodendrum* sporophylls do not take on identical positions at the beginning of their respective active phases (Fig. 2). The mean stigma angle (167.1°) in the natural population is close to the pollen deposition peak (170.1°). However, the mean anther angle (150.9°) at the beginning of the male phase is away from the pollen export peak (190.7°). In addition, while the stigma is stationary in its active phase ^16^, the anthers move at the rate of 1° per 0.36 hr (Supplementary Fig. 1). As a result of that pollination accuracy changes over time (Supplementary Fig. 3).

Anther movement during the active male phase results in a wide pollen export angle (Fig. 4A). In the counterfactual situation, if anthers were stationary, pollen would be exported from a restricted position and this would result in a narrow pollen export angle. Pollen export angle has a crucial role to play in pollination. In general, if pollination accuracy is maintained, we would expect that a narrow export angle would result in more efficient pollen export than would a wide export angle. The observed wide export angle leads to a lower deposition rate because pollen is exported from a broader zone, which includes the accuracy peak but is not exclusively restricted to it.

If a narrow export angle is advantageous, then why is the pollen export angle of *Clerodendrum* so wide (Fig. 4A)? Another way of asking this is why are anthers not stationary during the male phase? Further, why are anthers positioned at a non-optimal angle at the beginning of the male phase, when pollen load is high (Fig. 4B)? If variation in stigma angle is high in the population, then a wide pollen export angle might ensure pollen deposition at various positions. Moreover, there is an observed trade-off between pollen export and pollination accuracy (i.e., the deposition rate of exported pollen) (Fig. 4A) possibly because of the presence of pollen thieves in the visitor assemblage ^22,48^. During the male phase, the deposition rate of exported pollen gradually increases (Fig. 4A), while the amount of pollen available to be exported decreases (Fig. 4B). A resolution to this conundrum could come from the recognition that only a few pollen grains are sufficient for pollination ^49,50^, implying that there would be no point in flooding optimally placed stigmas with additional pollen. In general, optimal resource use would involve increasing quantity where quality is low and reducing quantity where quality is high. This may be what *Clerodendrum* flowers do to maximise male fitness. This strategy could result in the fertilisation of the greatest number of stigmas in the population, including those away from the optimal stigma position.

Contrary to the expectation that the natural population of anthers should be at the top of the delivery curve, at the beginning of the male phase anthers are far from the delivery peak (Fig. 4B). However realised male fitness is actually related to the number of delivered pollen, which in turn depends on not only on the delivery rate but also on the available pollen load (Fig. 4B). Additionally, estimating overall male fitness requires integrating pollen delivered over the entire male phase. How might overall fitness be related to the starting position of the anthers? One might naively expect that anthers should start out at the position associated with peak pollen delivery because pollen load is highest at the start of the male phase. Instead, theoretically starting at this position would lead to low pollen delivery, because of sexual interference ^7^. This means that pollen load should be high at angles where delivery rate is low, and as pollen load decreases, anther angle should change such that delivery rate increases. Multiplying these curves (Fig. 4C) and integrating over the duration of the male phase (Fig. 5) shows that, counter-intuitively, anthers should start away from the position of peak delivery in order to maximise total pollen delivery. This is precisely what is seen in the natural population, in which pollen can thus be delivered to flowers with a wide range of stigma angles (Fig. 5). Although this argument is based on empirical evidence from *Clerodendrum*, it is likely to hold true for all movement-assisted dichogamous species. This also implies that the long-standing notion that the style and stamen take identical positions in their active phases does not necessarily hold in movement-assisted dichogamy. Firstly, the stamens are not stationary. Secondly, even if one focusses on the initial position of stamen, this, too, deviates from the position of the style. Instead, the story is more complicated, and anthers use a sophisticated and dynamic strategy to maximise male fitness.

Pollination accuracy is a strong selection force in natural populations because it determines pollen deposition on conspecific stigmas ^11^. It is a form of stabilising selection and is expected to be high, especially in complete dichogamous species ^15,20,51^. However, the pollinator environment is rarely constant for a species on an evolutionary timescale. In fact, it varies considerably in space and time even within an ecological timescale. In such a situation, fluctuating selection ^52^ is the likely outcome. A generalist species that is visited by multiple pollinators whose relative abundance fluctuates over time and space is likely to experience different strengths of pollination accuracy selection. One reason is that accuracy selection strength could be pollinator-specific. For example, hawk moth pollinators might induce high accuracy selection strength, because they have a stereotypical and restricted approach angle to flowers: they always aligns themselves vertically in front of the flower and insert their proboscis from a distance. By contrast, the approach angle of butterflies is more variable, as they typically land on the inflorescence first and then insert their proboscis: this means that, depending on the landing position, the approach angle could be wide. In addition, the pollinating surface area of butterflies is large as their wings can carry pollen. On the contrary, in hawk moths, only the base of the proboscis (close to the head) can carry pollen. Due to these behavioural differences, pollination accuracy selection strength can differ from species to species, especially if they belong to different pollinator guilds. *Clerodendrum* is a generalist plant, visited by diverse groups of visitors, such as bees, butterflies, and hawk moths ^22,23^. Therefore it is important to investigate the effect of different strengths of pollination accuracy selection on anther movement dynamics. With increasing strength of pollination accuracy selection (Supplementary Fig. 2), the optimal anther starting position should come closer to the pollen delivery peak (Fig. 6C) as number of pollen delivered at initial anther angles decreases (Fig. 7).

**Fig. 7.**
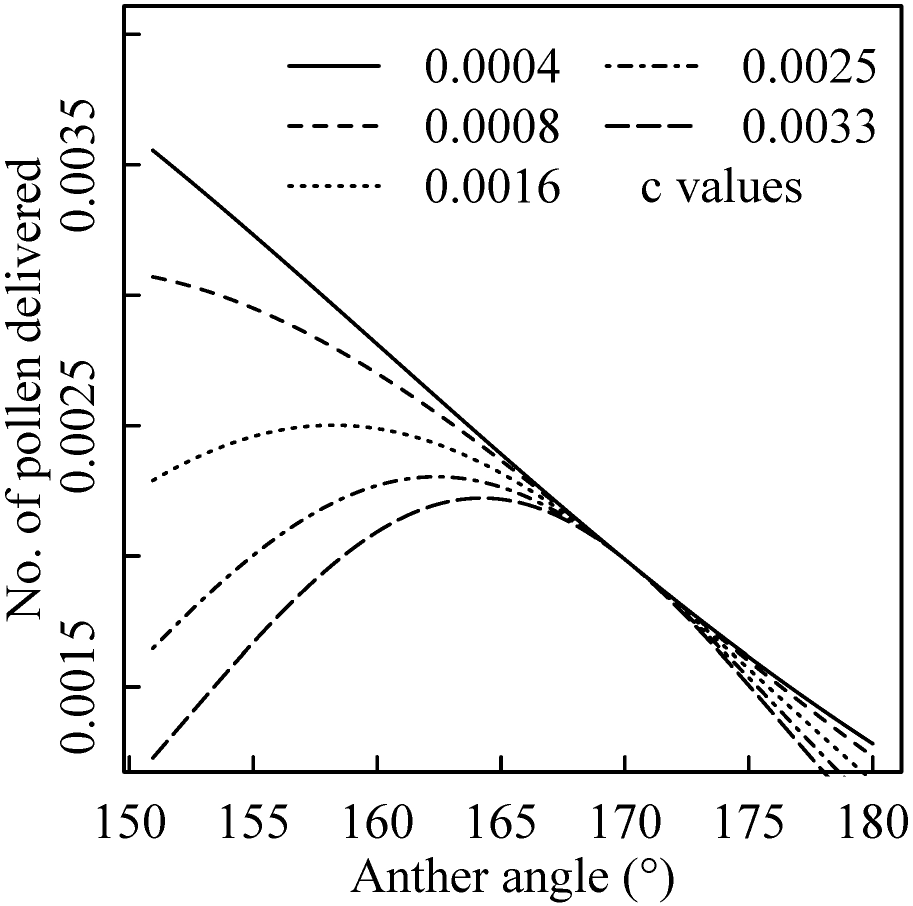
Number of pollen delivered at different anther angles at different pollination accuracy selection strength (parameter *c*).

Different pollinators are also likely to vary in their pollination efficiency. However, the model presented here suggests that this factor will not fundamentally change the optimum starting position of anthers; instead, it would alter the number of pollen delivered in the entire male phase (Fig. 6B). A further factor is visitation rate, which fluctuates greatly within and across populations. An increase in visitation rate would increase the total pollen delivered, and as well as shift the maximum pollen delivery peak towards the pollen export peak (Fig. 6A) as pollen gets quickly depleted at higher visitation rate (Fig. 8).

**Fig. 8.**
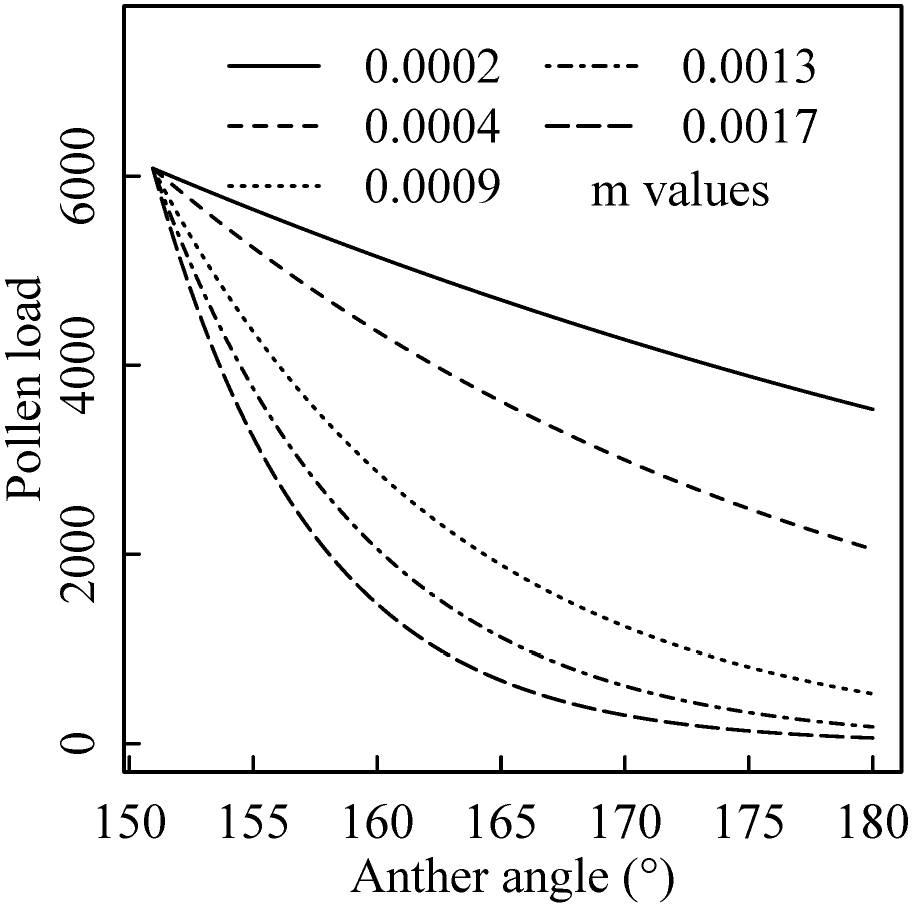
At different visitation rate (parameter *m*) change in pollen load as anthers move during the active phase.

For *Clerodendrum*, in which anthers move in their active phase, we have quantitatively shown that the starting position of anthers is close to the theoretical optimum. However, one might raise a more fundamental question about why anthers in their active phase are mobile at all. After all, a strategy of stable anther placement at the position that maximises delivery rate would generate the highest fitness. This question remains unresolved.

Overall, we conclude that *Clerodendrum* anthers are not in a stable position during their active stage: instead they move, and as a consequence pollination accuracy also changes over the male phase. Pollen export and pollination accuracy appear to trade off and their peaks do not align. Anthers start away from the position where pollen delivery rate is maximum. We demonstrate that overall male fitness, which depends on the total pollen delivered over the entire male phase, would in fact be lowest if anthers start at this position. Instead, starting away from the delivery peak means that pollen load is high where delivery is low, and pollen load is low where delivery rate is high. We argue that, because of sexual interference, this strategy in fact leads to the best outcome. In this way dynamic anthers maximise male fitness in a rather complicated way. Although our model was built to explain movement-assisted dichogamy specifically in *Clerodendrum*, a similar approach could be used to model movement-assisted dichogamy in general. In conclusion, by testing a variety of implicit assumptions that have been made about floral adaptations, our study reveals how ecological aspects of pollination can impact the intricacies of floral adaptation and evolution.

