## Supplemental Information for "Optimal inaccuracy: estimating male fitness in the movement-assisted dichogamous species *Clerodendrum infortunatum*"

The following Supplementary Information is available for this article:

**Supplementary Fig. 1** Anther movement with time in male phase of *Clerodendrum*.

**Supplementary Fig. 2** Pollen deposition rate at different strength of pollination accuracy selection.

**Supplementary Fig. 3** Positional mismatch between anther and stigma in their respective active phases.

**Supplementary Fig. 1** Anther movement with time in the male phase of *Clerodendrum*. Error bars represent 95% nonparametric confidence intervals. The fitted line ($y = 2.79x + 143.82$) is plotted, where $y$ = anther angle (°), $x$ = time (hr).

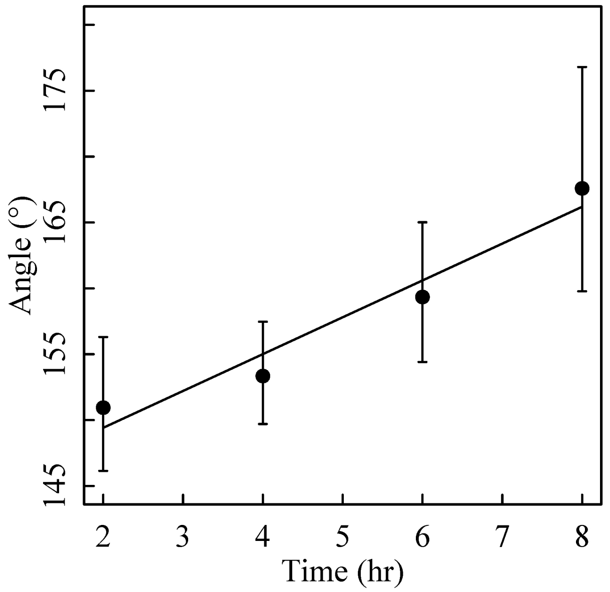

**Supplementary Fig. 2** Pollen deposition rate at different strength of pollination accuracy selection (parameter *c*). Note that pollen deposition rate is a very small number because it is standardised to a short time period (0.36 hr).

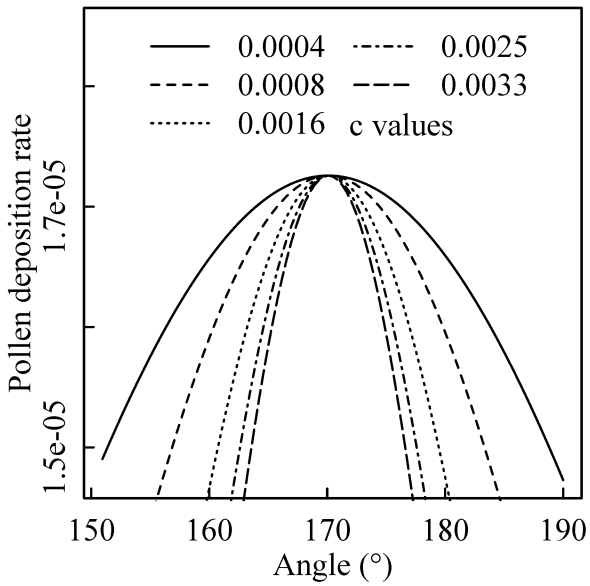

**Supplementary Fig. 3** Positional mismatch between anther and stigma in their respective active phases is measured in angles (°) and plotted against the time after anthesis in the male phase of the flower. Error bars are ±1 pooled standard error of mean.

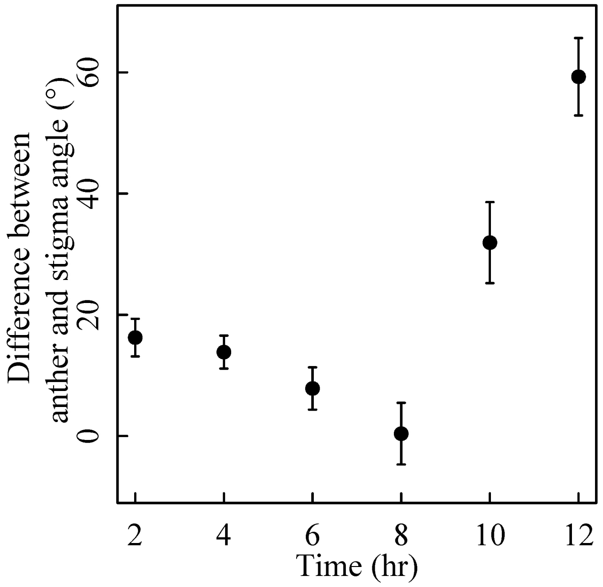
